# Placental malaria is associated with a TLR–Endothelin-3–oxidative damage response in human placenta tissues

**DOI:** 10.1101/2024.04.17.589949

**Authors:** Samuel Chenge, Melvin Mbalitsi, Harrison Ngure, Moses Obimbo, Mercy Singoei, Mourine Kangogo, Bernard N. Kanoi, Jesse Gitaka, Francis M. Kobia

## Abstract

Placental malaria, which is mainly caused by the sequestration of *Plasmodium falciparum*- infected erythrocytes in the placenta, is an important driver of poor pregnancy outcomes, including fetal growth restriction, preterm birth, and stillbirth. However, the mechanisms underlying its adverse outcomes are unclear. Mouse models have shown that placental malaria triggers a proinflammatory response in the placenta, which is accompanied by a fetal Toll-like receptor (TLR)4-mediated innate immune response associated with improved fetal outcomes. Here, we used hematoxylin and eosin staining to identify placental malaria positive and negative samples in our biobank of placentas donated by women living in a malaria-endemic region of Kenya and assessed the impact of placental malaria on the expression of TLRs, Endothelins, and oxidative damage. RT-qPCR analysis revealed that placental malaria was associated with an upregulation of TLR4, TLR7, and Endothelin-3. Moreover, immunohistochemistry showed that placental malaria was associated with elevated expression levels of the oxidative DNA damage marker, 8- hydroxy-2’-deoxyguanosine, while RT-qPCR revealed that this was accompanied by an upregulation of p21, an inhibitor of cell cycle progression and marker of cellular response to DNA damage. These findings allude to a novel mechanism of placental malaria pathogenesis driven by a TLR–Endothelin-3–oxidative DNA damage signaling axis.

## 1. Introduction

According to the World Health Organization, globally, there were about 249 million malaria cases and 608,000 malaria-associated deaths in 2022, with sub-Saharan Africa accounting for most of the cases and deaths [1]. Pregnant women have a higher susceptibility to malaria infection [2] and it is estimated that in 2022, there were about 12.7 million cases of malaria in pregnancy in sub-Saharan Africa [1]. Malaria in pregnancy is associated with several adverse outcomes in the mother, fetus, and neonate [3]. For the fetus, malaria in pregnancy severely worsens pregnancy outcomes and frequently leads to fetal growth restriction (including low birthweight, small for gestational age, and intrauterine growth restriction) and may result in preterm birth and stillbirth [3–5].

The adverse effects of malaria in pregnancy on the fetus are attributable to malaria infection of the placenta [1], leading to placental malaria. Placental malaria is characterized by the sequestration of *Plasmodium*-infected erythrocytes in placental intervillous spaces. This phenomenon is most frequently associated with *Plasmodium falciparum* (*P*. *falciparum*), the species associated with the most severe form of malaria [6]. The sequestration of *P. falciparum*- infected erythrocytes in the placenta is mediated by the interaction between variant surface chondroitin surface antigen 2, a *Plasmodium falciparum* protein expressed on the surface of infected erythrocytes [7], and chondroitin sulfate A on the surface of the syncytiotrophoblast [7], the placental epithelial cell layer that contacts maternal blood [8].

The adverse impacts of placental malaria on fetal well-being most likely result from the negative effects of placental malaria on placental health and function, since the vertical transmission of malaria to the fetus is rare [9]. Indeed, placental malaria is reported to induce placental inflammation [10,11] and placental histological changes [12], which may contribute to placental insufficiency and poor pregnancy outcomes. However, the mechanisms underlying placental malaria-driven placental pathobiology are not fully understood at the cellular and cell signaling levels.

Innate immune factors, such as Toll-like receptor (TLR)4, 7, and 9, which respond to infection by recognizing invading pathogens for clearance, are reported to respond to malaria infection [13], although their role in placental malaria is unclear. Although several studies have investigated maternal responses to placental malaria, few have studied how the fetus responds to parasite sequestration in the placenta. Nonetheless, a mouse model revealed that placental malaria triggers a TLR4-mediated innate immune reaction that adversely affects fetal outcomes, which is countered by a fetal innate immune reaction that led to better pregnancy outcomes [14]. This suggests the presence of TLR-mediated innate immune responses to placental malaria, although this has not been reported in the context of human placental malaria. Here, considering that mouse data show that TLR4 modulates endothelin-1 expression [15], malaria is inflammatory and oxidative [16], oxidative DNA damage upregulates TLR4 [17], and TLR signaling is thought to promote DNA repair [18], we used biobank placenta samples donated by women living in a malaria-endemic region of Kenya to examine the hypothesis that human placental malaria triggers a TLR–Endothelin–oxidative damage signaling response.

## 2. Materials and methods

### 2.1 The biobank

The study used biobank placenta samples donated by residents of Bungoma County, a malaria-endemic region of Western Kenya. The biobank was established by a previous prospective study as described recently [19]. This study did not require participant written informed consent since it was granted during biobanking [19]. Participants with a known record of sexually transmitted disease infection during pregnancy, those with pregnancy-associated noncommunicable diseases (preeclampsia and gestational diabetes) during the current pregnancy (at placenta donation), and those with twin pregnancies were excluded from analyses. Based on the underlying biobank data, malaria in pregnancy was defined as having at least one episode of hospital-diagnosed malaria during pregnancy. All data underlying the databank were deidentified. Equal numbers of male and female placentas were analyzed. The characteristics of the biobank’s placenta donors and samples are summarized in Table 1.

**Table 1.** Summary of placenta donors’ demographics.

| Group | Age<br>(range) | Gravidity<br>(range) | BW (g)<br>(range) | PW (g)<br>(range) | BW:PW<br>(range) |
| --- | --- | --- | --- | --- | --- |
| All donors | 24.7 years<br>(18–40) | 2.69<br>(1–7) | 3077.64<br>(1600–5000) | 479.04<br>(280.37–715) | 6.51<br>(3.2–10.2) |
| MiP | 24.4 years<br>(18–30) | 2.5<br>(1–7) | 2870.5<br>(1600–5000) **** | 464.3<br>(280.4–675) ** | 6.27<br>(3.2–10.2) * |
| NoMiP | 25 years<br>(18–40) | 2.9<br>(1–7) | 3272.3<br>(2000–4500) | 492.9<br>(342.3–715) | 6.73<br>(4.18–10) |

Analysis of the placenta donors' data revealed that maternal age and gravidity were not significantly different in the MiP (group with known history of malaria in pregnancy) vs the NoMiP (group without known MiP history) groups ( $P = 0.45$ and $0.12$ , respectively), whereas birthweight (BW), placental weight (PW), and BW:PW ratios were significantly lower in the MiP group when compared with the No MiP group ( $P = < 0.0001$ , $0.009$ , and $0.03$ , respectively). \*, \*\*, and \*\*\*\* indicate $P < 0.05$ , $0.005$ , and $0.0005$ , respectively.

### 2.2 Histological analysis

Formalin-fixed placenta tissues were embedded in paraffin blocks as previously described [20] using an automated tissue embedding system (MediMeas). H&E analysis was used to confirm placental malaria presence, which is indicated by the presence of infected erythrocytes in the placenta. Briefly, formalin-fixed paraffin-embedded samples were sectioned onto charged microscope slides (Dako, Cat No. K8020) at a 5-µm thickness, dried at 37 °C overnight on a slide warmer, dewaxed in xylene (Finar Chemicals, Cat No. 21940LC250), rehydrated by dipping across an alcohol gradient of absolute, 95%, 70%, and 50% ethanol (Scharlau, Cat No. ET00052500), and then in distilled water. They were then submerged in hematoxylin (Loba Chemie, Cat No. 04023) for seven minutes, rinsed with running water, and then destained through 10 dips in acid alcohol (1% hydrochloric acid in 70% ethanol). Next, they were submerged in eosin (Griffchem, Cat No. 45380) for 45 seconds followed by dehydration in 95% ethanol and absolute ethanol (five minutes each) and then cleared in xylene baths (10 minutes each) before being cover-slipped using dibutylphthalate polystyrene xylene mountant (Finar Chemicals, Cat No. 10525LM250). The slides were then examined under a microscope (Richter Optica UX1, M2 Scientifics), followed by imaging in ≥10 fields of view per slide at a 40X magnification using a Moticam microscope camera (Motic Scientific). Placental malaria was then diagnosed as described before [21], based on the presence of infected erythrocytes. Placental malaria-associated histopathological features were assessed by counting the number of syncytial knots as described previously [22], and measuring the fibrin-occupied placental areas using imageJ. These analyses were done in at least 10 fields of view per sample. The levels of placental malaria burden in the placental malaria-positive samples were determined by counting the number of identified infected erythrocytes in at least 10 fields of view per slide imaged at a magnification of 40X.

### 2.4 RNA extraction and reverse transcription quantitative PCR (RT-qPCR)

Total RNA was extracted from placenta tissue using a HigherPurity™ Tissue Total RNA Purification kit as per the manufacturer’s guidelines (Canvax, cat No. AN0152) and quantified using a NanoDrop Microvolume Spectrophotometer (ThermoFisher Scientific) following the manufacturer’s instructions. For each sample, 500 ng of RNA were retrotranscribed into cDNA using a LunaScript™ RT SuperMix cDNA Synthesis Kit (NEB, Cat. No. E3010L) using the manufacturer’s protocol. RT-qPCR analysis was done on a QuantStudio™ 5 Real-Time PCR System in a final volume of 20 µl containing 10 µl of GoTaq qPCR Master Mix (Promega, Cat No. PRA6001), 2 µl of the forward plus reverse primers (final primer concentration: 500 nM), 3 µl of nuclease free water (Promega, Cat No. P119E), and 5 µl of cDNA using the following program: 50 °C for two minutes, 95 °C for 10 minutes, followed by 40 cycles at 95 °C for 15 seconds and 60°C for 30 seconds. Relative gene expression was determined using the 2^-ΔΔct^ method [23], using β-actin as the reference gene. Primers were purchased from Macrogen and primer sequences are provided in Table 3.

### 2.5 *P. falciparum* detection PCR

The presence of *P. falciparum* in placenta samples was evaluated using One Taq® Quick-Load® 2X Master Mix with Standard Buffer (NEB, Cat No. M0486L) and the following thermocycler program: Initial denaturation at 95 °C for five minutes, followed by 35 cycles of denaturation at 95 °C for 30 seconds, annealing at 55 °C for 60 seconds, and extension at 72 °C for 75 seconds, and then a final extension at 72 °C for five minutes. Primer sequences are shown in Table 3. The PCR product was subjected to agarose (Sigma–Aldrich, Cat No. A9539) gel electrophoresis using 1X tris–borate–EDTA buffer alongside a 100 base pair ladder using SafeView™ Classic (Applied Biological Materials, Cat No. G108) nucleic acid stain and Gel Loading Dye, Purple (6X) (NEB, Cat No. B7024S). The bands were developed and imaged using a UVITEC Gel Documentation System (Cleaver Scientific).

### 2.6 Immunohistochemistry

The sections were deparaffinized by warming at 55 °C for 15 minutes followed by dipping in three xylene baths, about 10 dips each. They were then rehydrated and subjected to heat-induced epitope retrieval by boiling for 30 minutes in Citrate Buffer, pH 6.0 (Sigma–Aldrich, cat. No. C9999). They were then cooled to room temperature, rinsed with distilled water for five minutes and then blocked with 0.3% Triton-X in 1X phosphate-buffered saline (PBST). Next, they were blocked with 10% normal donkey serum (Abcam, cat. No. ab7475) in PBST for two hours followed by overnight incubation (4 °C) with anti-DNA/RNA damage antibody [15A3] (Abcam, cat. No. ab62623) at 1:2500 in blocking solution. Sections were then washed thrice (10 minutes each) using PBST and then incubated at room temperature for two hours with horseradish peroxidase-conjugated goat anti-mouse secondary antibody (Jackson ImmunoResearch, cat. No. 115-035- 003) at 1:5000 in blocking solution. The sections were then washed thrice (10 minutes each) using PBST followed by signal development using an ImmPACT® DAB Substrate Kit (Vector, cat. No. SK-4105) as per the manufacturer’s protocol. They were then dehydrated using 95%, 95%, 100%, and 100% ethanol (five minutes each), cleared by dipping in three xylene baths and then cover- slipped using a xylene-based mountant and allowed to dry. They were then examined under a light microscope and imaged at a magnification of 40X.

### 2.7 Data analyses

Statistical analyses were done using GraphPad Prism version 9. Data are presented as percentages, raw values, or mean ± standard deviation. Differences between two groups were compared using a t-test. Correlation analyses were done using nonparametric Spearman correlation analysis. For each placenta, the placental malaria burden in placentas was indicated by the total number of infected erythrocytes in the placenta section. The birthweight-to-placenta weight ratio was obtained by dividing the birthweight (grams) with the corresponding placenta’s weight (grams). The correlation between placental malaria status and birthweight, placental weight, and birthweight-to-placental weight ratio was assessed using GraphPad prism to examine the impact of placental malaria on fetal outcomes. *P* < 0.05 indicates statistically significant differences.

## 3. Results

### 3.1 Main characteristics of the placenta donors and donated placentas

All placenta donors were ≥18-years-old. The mean age, gravidity, birthweight, placental weight, and birthweight-to-placental weight ratio of the cohort of placenta donors were 24.7 years, 2.69, 3077.64 g, 470.04 g, and 6.51, respectively (Table 1). Grouping the placenta donors into those with a known history of malaria in pregnancy (MiP) and those without (NoMiP), revealed that in the MiP vs NoMiP groups, maternal age and gravidity were not significantly different (mean age: 24.4 [range: 18–30] vs 25 [range: 18–40] years, *P* = 0.45; mean gravidity: 2.5 [range: 1–7] vs 2.9 [range: 1–7), *P* = 0.12). However, in the MiP vs NoMiP groups, birthweight (mean: 2870.5 [range: 1600–5000] vs 3272.3 [range: 2000–4500] g, placental weight (mean: 464.3 [range: 280.4–675) vs 492.9 [range: 342.3–715] g, and birthweight-to-placental weight ratio (mean: 6.27 [range: 3.2– 10.2] vs 6.73 [range: 4.18–10], were significantly lower in the MiP group (*P* < 0.0001, = 0.009, and = 0.03, respectively). The presence of placental malaria was determined using H&E analysis as we described recently [19]. Of the placentas, 92 (51.4%) belonged to male babies, and eight (4.5%) were preterm (Table 2). All downstream analyses were done on placenta samples that were confirmed to be placental malaria-positive or placental malaria-negative using H&E staining.

**Table 2.** Main characteristics of the donated placentas.

|  |  |
| --- | --- |
| Placental malaria-positive placenta samples (infected erythrocytes present) | 92 (51.4%) |
| Placental malaria-negative placenta samples | 58 (32.4%) |
| Placentas with past placental malaria infection (hemozoin present) | 29 (16.2%) |
| Placenta samples from male fetuses | 92 (51.4%) |
| Placenta samples from female fetuses | 87 (48.6 %) |
| Placenta samples from pre-term deliveries | 8 (4.5%) |

The general characteristics of the placentas used in this study are summarized. Placental malaria status was determined using hematoxylin and eosin (H&E) staining of placental tissue. Placental malaria: observation of at least one infected erythrocyte [46]. Past placental malaria: absence of infected erythrocytes coupled with hemozoin in fibrin and pigmented macrophages [21]. Chronic placental malaria: detection of infected erythrocytes, as well as hemozoin in fibrin and pigmented macrophages (past placental malaria) [47].

**Table 3.**
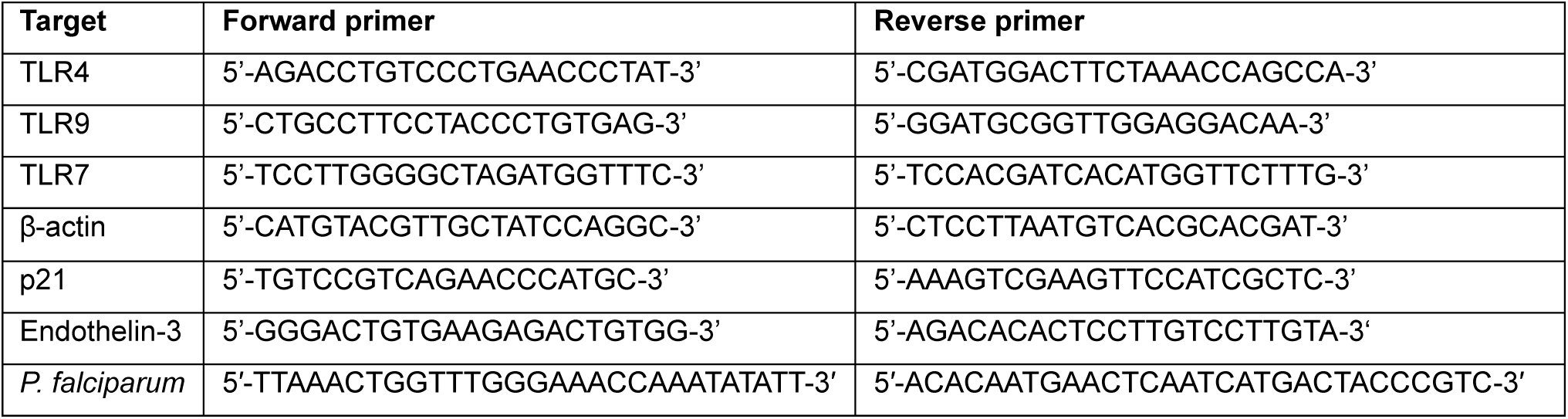
List of primers used in the study.

### 3.2 Placental malaria correlates negatively with birthweight and birthweight-to-placenta ratio

Representative images of placental malaria-negative and placental malaria-positive samples are shown in Figure 1A–B, and the presence of *P*. *falciparum* in placental malaria-positive tissues was confirmed using PCR (Figure 1C). Analysis of the data underlying the placental biobank revealed that when compared with the placental malaria-negative group, birthweight was significantly lower in the placental malaria-positive group, which had more low birthweight cases (Figure 1A, *P* = 0.03, low birthweight: <2500 g), but placental weight did not differ between the two groups (Figure 2B, *P* = 0.8). However, the birthweight-to-placental weight ratio was lower in the placental malaria-positive group, although the difference did not reach statistical significance (Figure 1F, *P* = 0.1). Next, we used the H&E images to determine the proportion (%) of infected erythrocytes (IEs) in each placenta sample, and then used the obtained values to assess the correlation between the placental malaria burden and birthweight, placental weight, and the birthweight-to-placental weight ratio, i.e., the fetal weight obtained per gram of the placenta, which is an indicator of placental efficiency, with higher birthweight-to-placental weight ratios indicating greater placental efficiency [24]. This analysis revealed negative correlation between IEs (proportion of infected erythrocytes [%]) and birthweight (correlation coefficient [rs]: −0.23, *P* < 0.005, 95% confidence interval [CI]: −0.37 – −0.08), and IEs (%) and birthweight-to-placental weight ratio (rs: −0.20, *P* = 0.007, 95% CI: −0.35 – −0.05). Expectedly, placenta weight had a positive correlation with birthweight (rs: 0.28, *P* < 0.001, 95% CI: 0.13 – 0.41) and a negative correlation with birthweight-to-placental weight ratio (rs: −0.42, *P* < 0.001, 95% CI: −0.54 – −0.28). However, placental malaria burden did not exhibit correlation with placenta weight (rs: 0.01, *P* = 0.93, 95% CI: −0.16 – 0.15). Taken together, these findings indicate that placental malaria impairs placenta function, leading to low birthweight via reduced placenta efficiency, as indicated by the negative correlation between placental malaria burden and the birthweight-to-placental weight ratios of placental malaria-exposed neonates.

**Figure 1.**
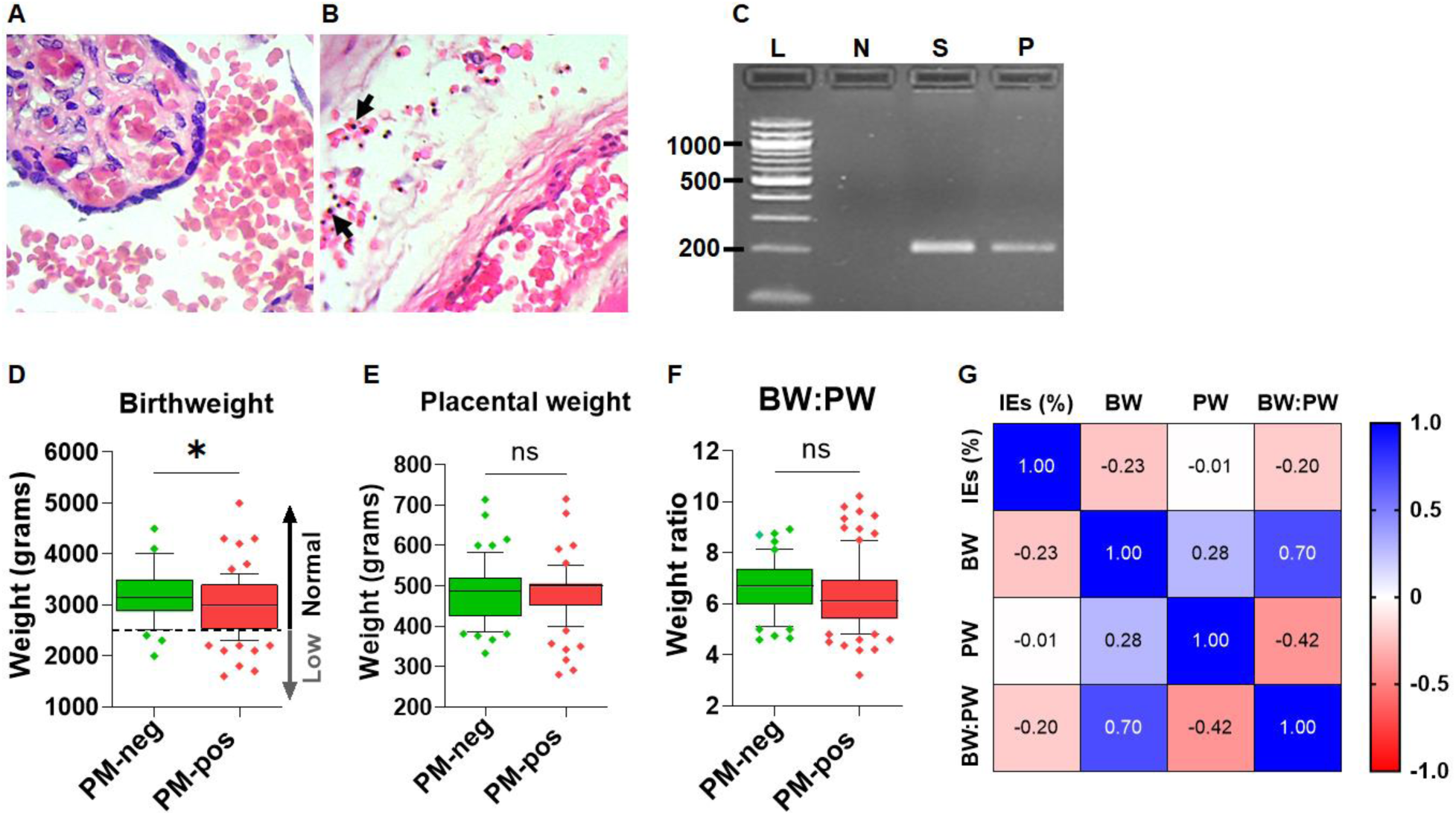
(A–B) Representative hematoxylin and eosin images of placental malaria (PM)-negative. (A) and PM-positive samples (B). When compared with a PM-negative sample, the positive sample has malaria-infected erythrocytes (black arrowheads) in the placental intervillous space. (C) PCR confirmed the presence *of P. falciparum* in the PM-positive sample in B. L: 100 base pair ladder, N: PM-negative sample, S: PM-positive sample in B (histology), P: positive control (*P. falciparum* strain 3D7 genomic DNA). Expected PCR band size: 205 base pairs. (D–F) PM was associated with lower birthweight (BW) (D, *P* = 0.03) and lower birthweight-to-placental weight (BW:PW) ratio, although the difference did not reach statistical significance (F, *P* = 0.1), but not with lower placental weight (PW) (E, *P* = 0.8). In D–F, whiskers are drawn from the 10^th^ to 90^th^ percentile. (G) A correlation matrix shows that the proportion of infected erythrocytes, IEs (%), in the placenta correlated negatively with BW (*P* < 0.005) and the BW:PW ratio (*P* = 0.007), but it did not correlate with PW (*P* = 0.94). PW correlated positively with BW and negatively with the BW:PW ratio (both *P* < 0.001).

**Figure 2.**
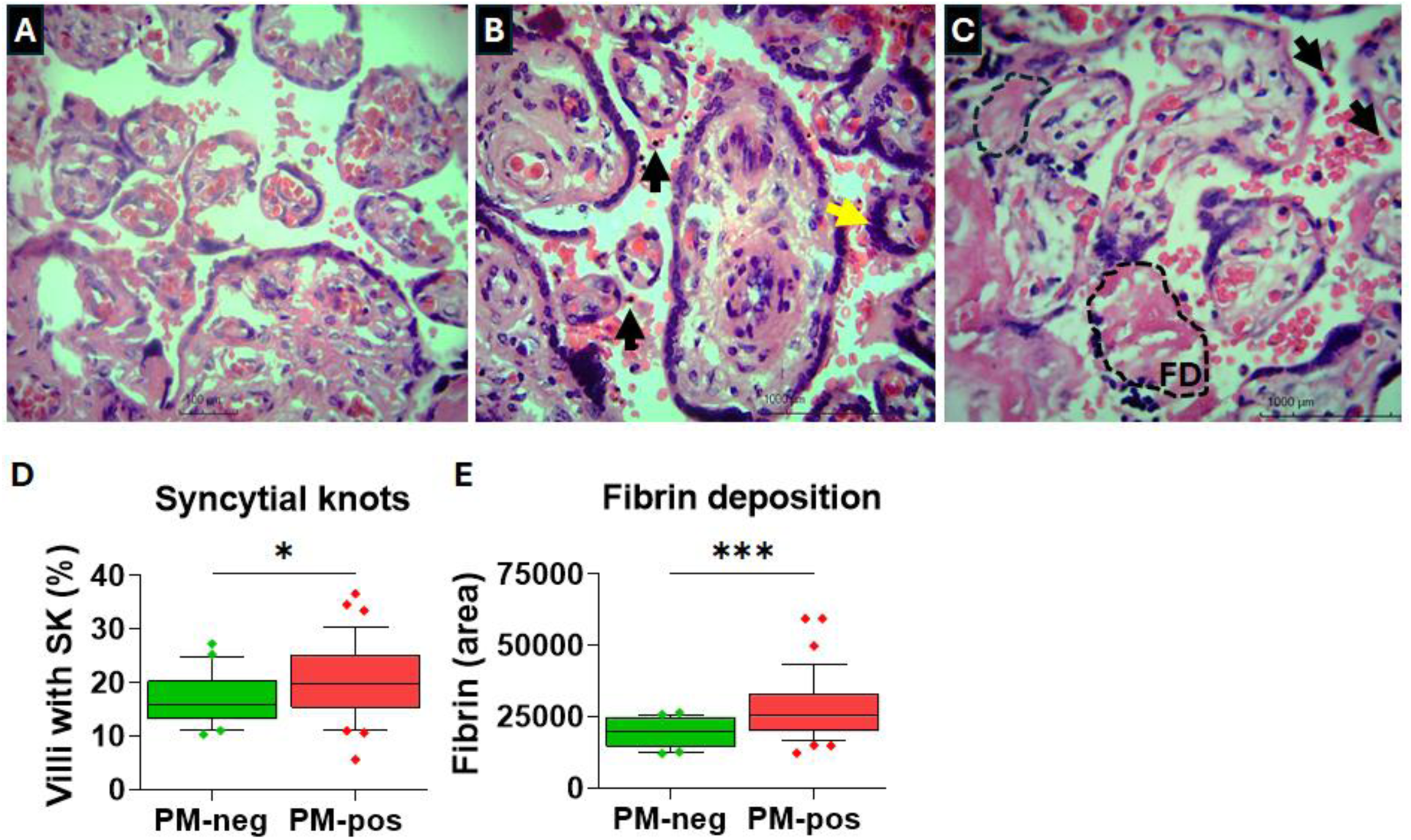
(A–C) In the samples from the biobank underlying our study, when compared with placental malaria (PM)-negative samples (A), PM was associated with significantly higher rates of syncytial knots (B, yellow arrowhead) and placental fibrin deposits (C; FD, marked with broken line). Black arrowheads indicate infected erythrocytes. (D–E) Quantification revealed that the levels of syncytial knots (SK [D], n = 21 and 38 for PM-neg and PM-pos, respectively; *P* = 0.047) and the area of placenta intervillous space containing fibrin (E, n = 26 and 38 for PM-neg and PM- pos, respectively, *P* < 0.0005) were significantly higher in the PM-positive (PM-pos) samples than in the PM-negative (PM-neg) samples. Whiskers are drawn from the 10^th^ to 90^th^ percentile.

### 3.3 Placental malaria markedly alters placental histological features

Next, we assessed the impact of placental malaria on syncytial knotting and fibrin deposition in our placenta samples. This analysis revealed that when compared with placental malaria-negative samples (A), placental malaria-positive samples had more syncytial knots (B, yellow arrowhead) and greater fibrin-occupied placental area (C, FD [fibrin deposit] in the broken line-demarcated area). Quantification analyses revealed that when compared with placental malaria-negative samples, the levels of these histological features were significantly higher in placental malaria-positive samples (D, SK [syncytial knots], *P* = 0.047 and E, fibrin area, *P* < 0.0005). These observations indicate that in our study cohort, placental malaria may have adversely affected fetal outcomes by at least in part, altering normal placental histological features.

### 3.4 Placental malaria is associated with an upregulation of TLR4, TLR7, and Endothelin 3

We then sought to determine if placental malaria altered the expression of TLRs. To this end, we focused on TLR4, TLR7, and TLR9, which have been associated with response to malaria infection in mice [25,26], and with mouse placental malaria in the case of TLR4 [27], although this has not been reported in human placental malaria. To evaluate the effect of placental malaria on these innate immune system receptors, we assessed their expression levels using RT-qPCR. The analysis revealed that when compared with placental malaria-negative controls, placental malaria-positive samples expressed significantly higher levels of TLR4 and TLR7, but not TLR9 (Figure 3A–C, *P* = 0.002, 0.03, and 0.59, respectively). This is consistent with mouse data showing that placental malaria upregulates TLR4-mediated expression of endothelin-1 [15]. We therefore wondered if human placental malaria alters the expression of Endothelin genes. RT-qPCR analysis of Endothelin-1 and -3 gene expression revealed that only Endothelin-3 was detectable in our placenta samples and that when compared with placental malaria-negative placentas, placental malaria-positive samples had significantly higher Endothelin-3 levels (Figure 3D, *P* = 0.004), indicating the presence of a TLR–Endothelin signaling axis in response to human placental malaria.

**Figure 3.**
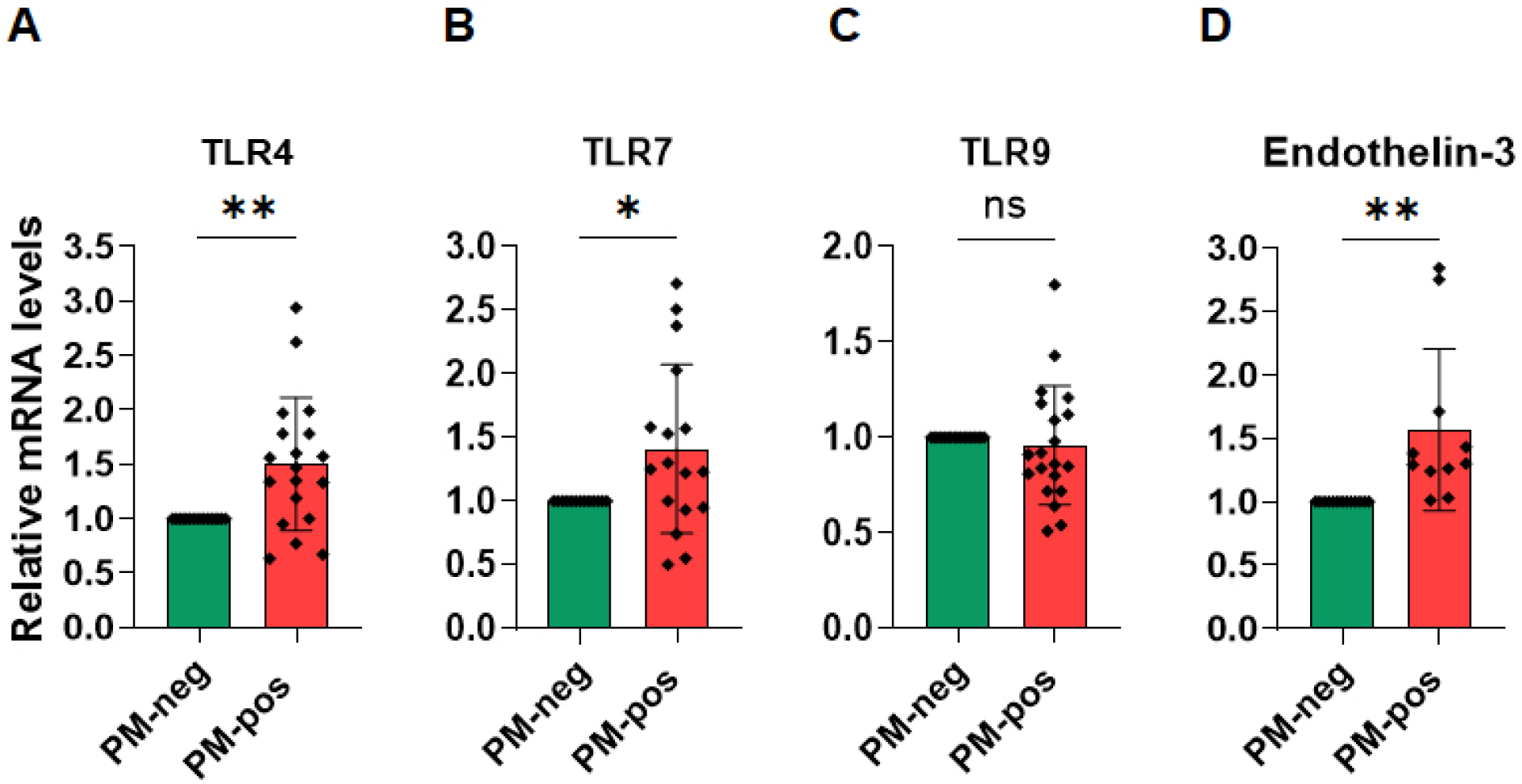
Placental malaria is associated with the upregulation of TLR4, TLR7, and Endothelin-3. (A–C) When compared with placental malaria (PM)-negative samples (PM-neg), PM-positive (PM-pos) samples expressed significantly higher levels of TLR4 and TLR7, but not TLR9 (*P* = 0.002, 0.03, and 0.59, respectively). (D) PM-positive samples also expressed higher levels of Endothelin-3 (*P* = 0.004).

### 3.5 Placental malaria is associated with high oxidative DNA damage

Since TLR4 was upregulated in placental malaria-positive placenta samples, we wondered whether placental malaria-driven TLR4 activation is associated with a dysregulation of other signaling processes that may underlie or contribute to placental malaria-mediated placental pathobiology. Because malaria is known to be strongly inflammatory and oxidative, which may drive host tissue damage [16], and because oxidative DNA damage is associated with TLR4 upregulation [17] and that TLR signaling is thought to promote DNA repair [18], we wondered if the TLR4 upregulation in the placental malaria-positive samples might be associated with placental oxidative DNA damage. To assess this possibility, we used immunohistochemistry to assess the levels of 8-hydroxy-2’-deoxyguanosine, a marker of oxidative DNA stress [28]. This analysis revealed that when compared with placental malaria-negative samples, placental malaria-positive samples express markedly higher levels of 8-hydroxy-2’-deoxyguanosine (Figure 4A). Staining the same samples with the secondary antibody only (without the primary antibody) confirmed signal specificity (Figure 4B). To further assess the effect of placental malaria on oxidative stress, we used RT-qPCR to examine the level of p21, a mediator of cell cycle arrest and indicator of cellular response to DNA damage [29]. This revealed that when compared with placental malaria-negative samples, placental malaria-positive samples had significantly higher levels of p21 (Figure 4C, *P* = 0.02). Taken together, these data indicate that placental malaria triggers markedly high levels of placental oxidative DNA stress, placenta tissue damage, and cellular response to DNA stress, which may contribute to placental malaria pathobiology, and that in response, TLR signaling may at least in part, be upregulated to counter these adverse effects through the promotion of DNA repair.

**Figure 4.**
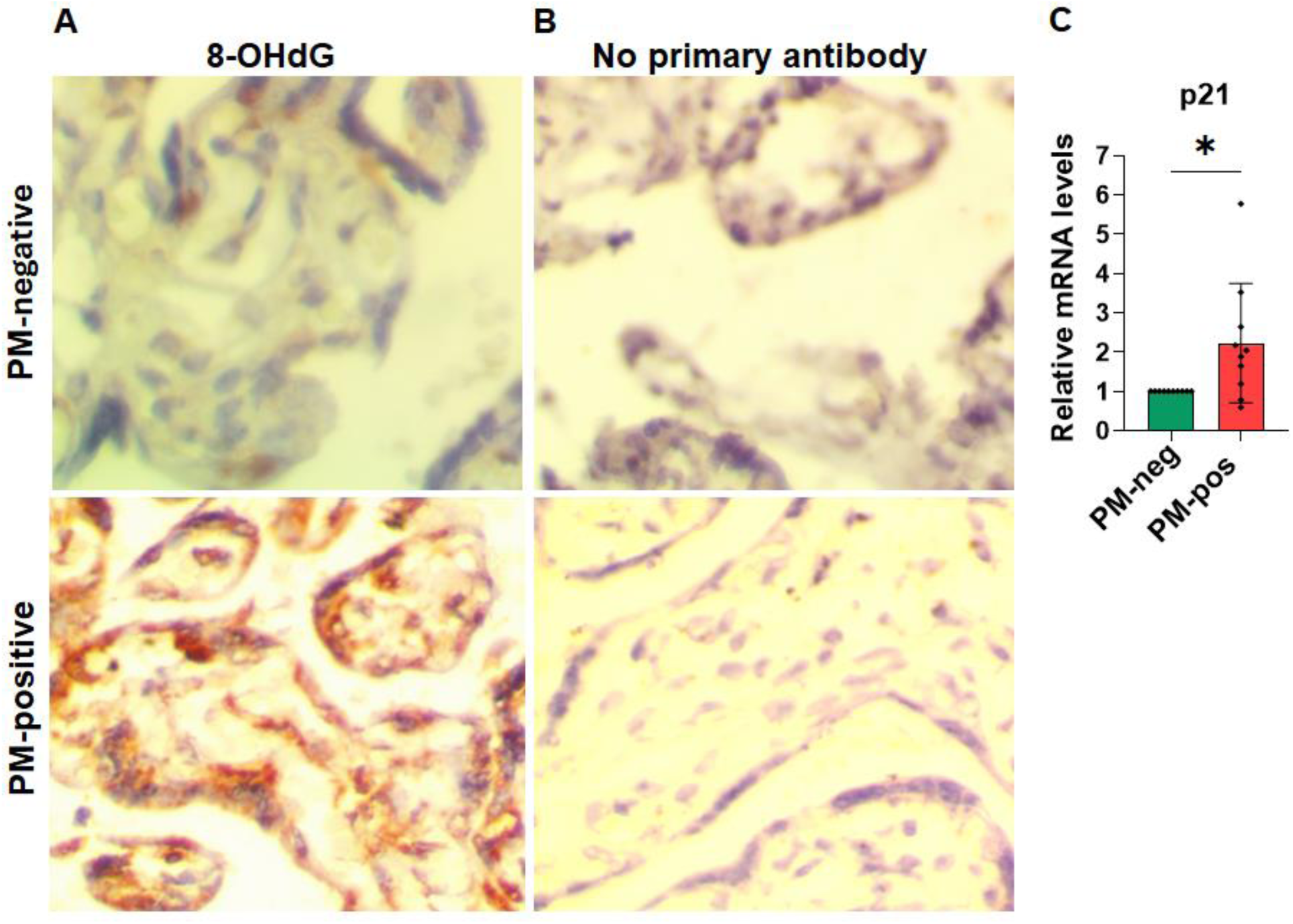
Analysis of oxidative DNA damage in placental malaria (PM)-negative vs PM-positive samples. (A–B) Immunohistochemistry revealed that when compared with PM-negative samples, PM-positive tissues had markedly higher levels of 8-hydroxy-2’-deoxyguanosine (8-OHdG), an indicator of oxidative damage. Staining the same samples with the secondary antibody only (B) confirmed signal specificity for this marker. (C) RT-qPCR showed that PM-positive samples express significantly higher levels of p21 (*P* = 0.02).

## 4. Discussion

Malaria in pregnancy often results in placental malaria, where erythrocytes that are infected with *P. falciparum*, the parasite that most frequently causes placental malaria, sequester in placental intervillous spaces [7]. Placental malaria may cause various adverse fetal outcomes, including stillbirth, preterm birth, and fetal growth restriction [3–5], and because *P*. *falciparum* rarely undergoes vertical transmission [9], these effects are likely caused by placental malaria-driven pathobiological effects in the placenta, which may impair placental function. However, although studies have implicated effects like inflammation and histological changes in placental malaria pathogenesis, the mechanisms underlying the adverse effects of human placental malaria on the placenta are unclear. Moreover, because many malaria in pregnancy cases in malaria-endemic regions are asymptomatic [30,31] and the placenta is inaccessible during pregnancy, there are no ways of detecting and intervening against placental malaria during pregnancy. Thus, there is an urgent need to better understand the placental pathobiology of placental malaria to guide the development of effective diagnostic and therapeutic tools.

Mouse models indicate that placental malaria triggers innate immune responses (mainly via TLR4) that are associated with poor fetal outcomes and that TLR4-mediated fetal responses to placental malaria cause improved outcomes [14]. However, the effect of human placental malaria on TLR-mediated immunity in the placenta has not been examined. In this study, we leveraged our well-characterized biobank of placenta samples from a malaria endemic region of Kenya (Table 1) and found that in our study cohort, placental malaria burden had a significant negative correlation with birthweight and birthweight-to-placental weight ratio, that it was associated with significantly higher placental histological lesions , higher levels of TLR4 and Endothelin-3 expression, and enhanced oxidative DNA damage when compared with samples without placental malaria.

Consistent with previous findings implicating placental malaria in fetal growth restriction [32], we observed that in our study cohort, relative to the placental malaria-negative cases, placental malaria was associated with low birthweight. Moreover, we observed that placental malaria was associated with lower birthweight-to-placental weight ratios, an indicator of placental efficiency in which higher ratios indicate higher nutrient transfer for every gram of the placenta and vice versa [24], but not with lower placental weight (Figure 1D–F), an observation that to our knowledge, has not been previously reported in human placental malaria. Interestingly, our analyses also indicate that the placental malaria burden (percentage of infected erythrocytes in a sample’s intervillous spaces) correlates negatively with birthweight but not with placental weight. Taken together, these observations indicate that placental malaria contributes to fetal growth restriction primarily by impairing placental function, and not via placental growth inhibition, although the precise mechanisms remain unclear. This possibility is crucial considering that women in malaria-endemic regions may experience multiple malaria reinfections throughout pregnancy. Therefore, further studies, such as using *in vitro* and organoid systems, are needed to comprehensively investigate this possibility.

Our analyses revealed that placental malaria-positive samples had markedly higher levels of fibrin deposition and syncytial knotting than placental malaria-negative samples, which is in line with earlier findings [33]. These changes, which indicate placental injury, and have been associated with placental malperfusion and poor fetal outcomes, including fetal growth restriction [34,35], may contribute to the low birthweight observed in our placental malaria cohort. However, studies are needed to establish the mechanisms by which placental malaria alters placental histological features, how these changes correlate with fetal outcomes and postnatal wellbeing, and whether they can predict fetal wellbeing in postnatal life.

TLRs are key innate immunity factors that sense host invasion by pathogens and activate host immune defenses [36]. Mouse models of malaria indicate that TLR4, TLR7, and TLR9 are involved in detecting and responding to malaria infection [25,26]. Moreover, mouse models indicate that at the fetal–maternal interface, placental malaria activates TLR4-mediated immune responses that drive poor fetal outcomes, and that fetal TLR4-mediated counterresponses improve pregnancy outcomes [14,27,37]. However, this observation has not been previously made in human placental malaria. Here, we observed that the expression levels of TLR4 and TLR7, but not TLR9, were significantly upregulated in placental malaria-positive samples, indicating that placental infection triggers an innate immune reaction and that it is mainly driven by TLR4 and TLR7, although the status of other TLRs during placental malaria warrants investigation. Considering that TLRs are important drivers of inflammation [38], which is implicated in placental malaria pathogenesis [39], taken together with the observed changes in placental histological features, our findings indicate for the first time, that TLR-mediated responses to placental malaria may contribute to local placental inflammation, which may underlie the observed placental malaria-associated low birthweight and placental weight, although the precise mechanisms remain unclear. Mouse data show that placental malaria-driven TLR4 expression drives placental endothelin-1 expression [15], and for the first time, our findings show that placental malaria is also associated with the upregulation of Endothelin-3. The Endothelin ligands 1, 2, and 3 are a family of vasoactive factors that influence a range of cellular processes, such as vascular remodeling and angiogenesis [40]. Moreover, Endothelin-3 is reported to have anti-inflammatory effects [41,42], suggesting that its upregulation in the context of placental malaria-mediated TLR4 upregulation is a mechanism of countering TLR4-dependent placenta inflammation. Collectively, these observations indicate that human placental malaria may activate a TLR4–Endothelin-3 signaling axis, but further studies are needed to test this hypothesis and to determine its implications in placental malaria pathobiology and fetal outcomes.

Based on reports that malaria is strongly oxidative [16], oxidative stress causes tissue damage [43], oxidative DNA damage upregulates TLR4 [17], and that the TLR pathway might promote DNA repair [18], we reasoned that our observation of TLR4 and TLR7 upregulation in placental malaria-positive samples might be accompanied by placental oxidative DNA damage. This hypothesis was confirmed by our immunohistochemistry data, which showed that 8-hydroxy-2’- deoxyguanosine, a marker of oxidative DNA stress [28], was markedly upregulated in placental malaria samples. Moreover, gene expression analysis revealed that these events were accompanied by a significant upregulation of p21, a cell cycle inhibitor and marker of cellular response to DNA damage [29]. These observations align with previous findings that in a mouse model, placental malaria is associated with placental oxidative damage [44]. Furthermore, p21 upregulation in the placenta may arrest the cell cycle to allow for oxidative damage resolution, which may contribute to the low placental weight observed in our cohort, but this possibility requires further investigation. Together, our data suggest the presence of a previously unknown TLR–Endothelin–oxidative damage axis in human placental malaria.

Finally, our data also show that in the placental malaria-negative and placental malaria-positive groups, the analyzed parameters varied across samples, which may reflect inter-individual biological variability [45]. In the future, our observations will be validated experimentally, such as by challenging cultured primary human trophoblasts with *P. falciparum* for detailed mechanistic interrogation.

## 5. Conclusion

Despite its heavy burden and adverse effects on maternal and fetal outcomes, the mechanisms underlying the placental pathobiology of placental malaria are unclear. Considering that malaria is rarely transmitted to the fetus [9], the adverse fetal outcomes of malaria in pregnancy are mainly driven by events that disrupt placenta physiology and function. Importantly, because of placental inaccessibility, placental malaria can only be confirmed via postnatal placental histopathology. These challenges highlight the urgent need to better understand the mechanisms underlying the placental pathobiology of placental malaria, which may inform the development of sensitive tools for diagnosing placental malaria during pregnancy as well as effective therapeutic interventions. Our findings that placental malaria may drive TLR-mediated responses in the placenta raise the possibility that modulating innate responses to placental malaria may improve fetal outcomes, as we previously discussed [13]. Moreover, our identification of an axis involving TLRs, Endothelins, and oxidative DNA damage during placental malaria (Figure 5) highlights processes with the potential for intervention against human placental malaria. However, further studies, such as using primary human trophoblasts, human placental organoids, or human placental *ex vivo* systems are needed to validate our observations. Such approaches can investigate the mechanisms of placental malaria pathobiology more rigorously than can be done using term placentas.

**Figure 5.**
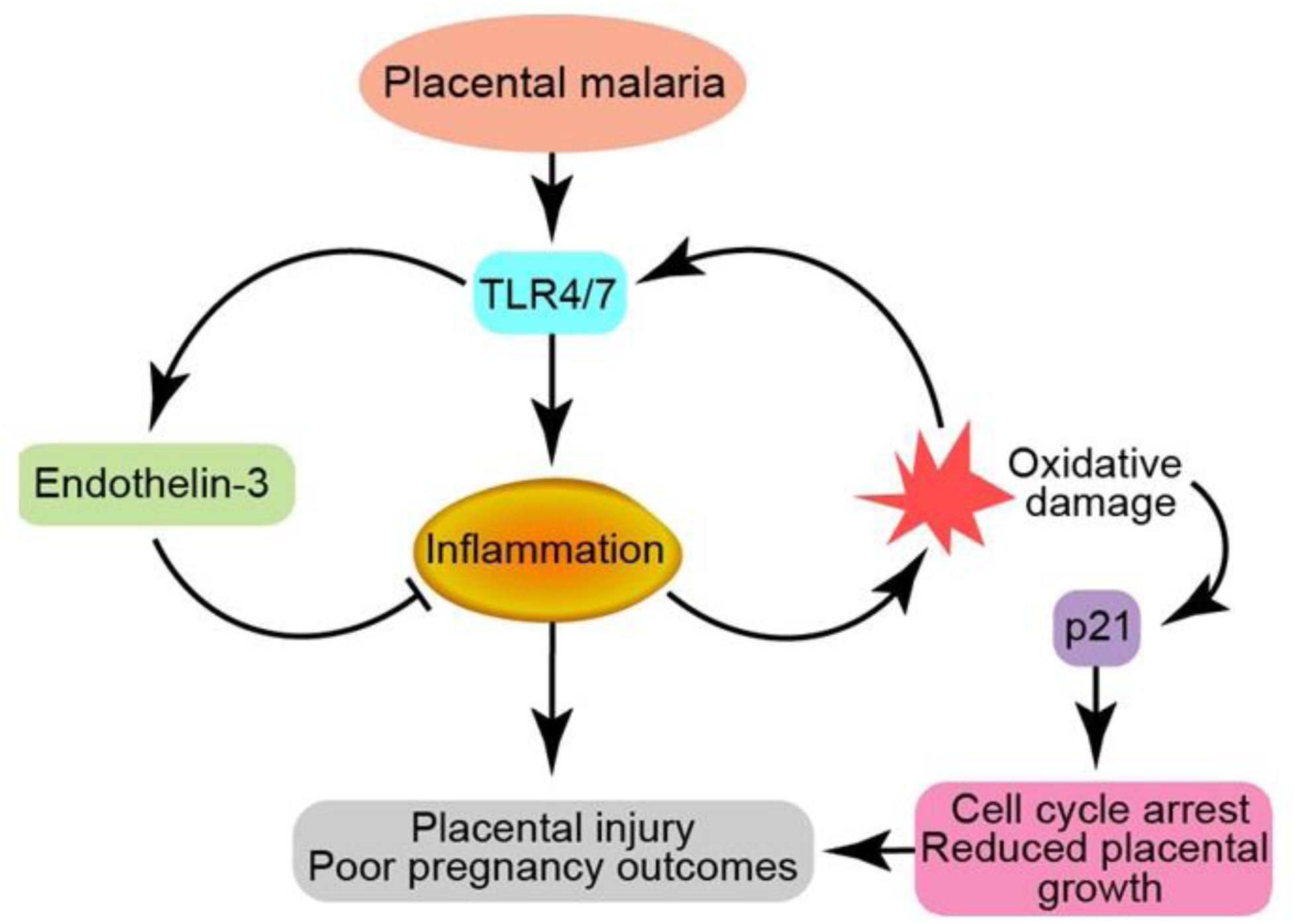
Schematic summary of the hypothesized TLR–Endothelin-3–oxidative stress axis in human placental malaria. Further investigation is needed to validate this axis and determine its potential for intervention against placental malaria.

## Acknowledgments

We thank our placenta donors and the staff at Webuye County and Mary Help of The Sick (Thika) Hospitals for their generous support. We thank the staff of the histopathology and anatomy departments at the University of Nairobi and Mount Kenya University for their help with histopathology. We are grateful to Prof. Walter Jaoko, Prof. Omu Anzala, and Dr. Daniel Muema of KAVI–ICR for kindly sharing laboratory space and resources. We are thankful to Prof. Roger Smith and Dr. Kaushik Maiti, Mothers and Babies Research Centre, Hunter Medical Research Institute, Newcastle, NSW – Australia, for sharing antibodies and other resources.

## Ethics statement

This study was approved by Mount Kenya University’s ethics review committee (approval number 2516).

## Data accessibility

The data underlying this study are contained in the article.

